# A Neurodevelopment-Inspired Deep Spiking Neural Network for Auditory Spatial Attention Detection using Single Trial EEG

**DOI:** 10.1101/2025.11.17.688781

**Authors:** Faramarz Faghihi, Ahmed Moustafa

**Affiliations:** Department of Medical Physiology, Division of Heart & Lungs, University Medical Center Utrecht, Utrecht, The Netherlands; School of Psychology, Faculty of Society and Design, Bond University, Gold Coast, Queensland, Australia; Department of Human Anatomy and Physiology, the Faculty of Health Sciences, University of Johannesburg, South Africa

**Author notes:** **Corresponding author:** Ahmed A. Moustafa.

**Keywords:** Deep learning, spiking neural networks, neurodevelopment, learning rule, sparse coding, EEG, auditory attention, machine learning, Self-Organizing, synaptic synchronization, learning from small datasets

## Abstract

This study presents a deep probabilistic spiking neural network designed to extract discriminative spatiotemporal features from EEG signals associated with rightward and leftward auditory attention. The network self-organizes its synaptic weights and inter-layer connectivity through a biologically inspired learning rule shaped by probabilistic feedback inhibition and spike-based synchronization dynamics. To characterize the model’s behavior and identify optimal operating conditions, we systematically examined the effects of key parameters including inhibition strength, synchronization parameters, and preprocessing thresholds on network stability and classification performance. Simulation results show that feedback inhibition is essential for preventing uncontrolled synaptic growth, maintaining sparse connectivity, and enabling stable learning. Strong inhibition suppresses connectivity and weight development, whereas weak inhibition leads to excessive synaptic expansion. An intermediate inhibition regime achieves a balance between adaptability and stability, resulting in the most effective feature extraction. Output-layer neurons exhibited heterogeneous activation dynamics, reflecting diverse encoding of EEG input patterns. The excitatory firing probability was tightly modulated by the inhibition parameter, confirming its central role in shaping network responses. Synchronization parameters further influenced synaptic dynamics, producing nonlinear effects on spike coordination and weight evolution. Classification accuracy peaked at intermediate synchronization levels, revealing an optimal regime where inhibition and synchronization jointly support efficient classification of rightward and leftward EEG data. Additional analyses demonstrated that synaptic pruning substantially improves accuracy across all inhibition levels and that the model performs best when input spikes are generated using a preprocessing threshold of 0.3. Moreover, experimental results demonstrate that the developed neural network model achieves an average accuracy of 90% while utilizing only 10% of the available EEG data. Overall, the findings show that the proposed spiking neural network achieves its highest classification performance under moderate inhibition, intermediate synchronization, sparse connectivity, and appropriate pruning. These results highlight the promise of biologically inspired spiking architectures for decoding EEG-based attentional states, particularly in settings with limited training data and strong temporal structure.

## Introduction

Electroencephalography (EEG) is a widely used noninvasive neurophysiological technique that captures voltage fluctuations generated by postsynaptic potentials in cortical neuronal populations. Owing to its millisecond-scale temporal resolution, EEG is ideally suited for examining the fast neural dynamics underlying perception, attention, memory, and decision-making. In cognitive neuroscience, EEG provides access to neural oscillations and event-related potentials (ERPs), offering insights into the temporal organization of brain activity and the mechanisms supporting information processing. Advances in computational modeling have further enabled the decoding of complex cognitive states from EEG signals, expanding the scope of research on brain function and cognition [Huang & Wang, 2021].

Machine learning (ML) has become a central tool for analyzing EEG data in cognitive and clinical neuroscience. Numerous studies have demonstrated the ability of ML models to extract discriminative spatiotemporal patterns related to attention, perception, working memory, and other cognitive operations [Hosseini, 2020; Aggarwal, 2022; Zeng, 2022; Saeidi, 2021; Ghosh, 2015; Delgado, 2025; Wang, 2025; Leung, 2025; Qi & Fengxue, 2025]. Traditional approaches employ regression, classification, and clustering algorithms, while more recent deep learning (DL) methods automatically learn hierarchical representations from raw or minimally processed EEG data [Lotte, 2007; Luján, 2021; Subasi, 2005; Hossain, 2023; Xu, 2021]. These developments have notably improved decoding performance and opened new avenues for cognitive monitoring and brain–computer interface (BCI) applications.

Despite their success, deep learning models are based on artificial neural networks (ANNs) composed of continuous-activation units whose computational mechanisms differ markedly from biological neurons. DL models typically require large datasets to avoid overfitting and rely on computationally expensive optimization procedures, such as backpropagation, that lack biological plausibility [LeCun, 2015; Mathew, 2020; Shinde, 2018; Panahi, 2025; Kang, 2021; Shrestha, 2019]. In parallel, neurophysiological studies have shown that auditory attention can be decoded from EEG, including the detection of spatial focus based on alpha-band activity [An, 2020; Wang, 2023; Lan, 2025; Deng, 2020]. Convolutional neural networks (CNNs) and frequency-domain EEG analyses have been successfully applied to classify the direction of auditory attention [Vandecappelle, 2021; Jiang, 2022; Mahjoory, 2024]. However, EEG signals are nonstationary, nonlinear time series, suggesting that models that explicitly exploit temporal dynamics such as spiking neural networks (SNNs) may offer distinct advantages.

Spiking neural networks are biologically inspired learning systems in which information is encoded by sparse, temporally precise spike trains rather than continuous activations [Tavanaei, 2019; Maris Ferreira, 2025]. Their sparse communication makes SNNs more energy-efficient than ANNs, an appealing property for portable or neuromorphic devices. Because SNNs operate on spike timing and firing rates, they naturally capture spatiotemporal structure in dynamic signals such as EEG [Yamazaki, 2022; Petro, 2019; Wu, 2018; Kugele, 2020]. These networks have been used to simulate neural activity at both single-neuron and population levels, supporting studies of cognitive processes [Fang, 2010]. SNNs have also demonstrated strong performance in applications including visual and auditory processing, speech recognition, medical diagnosis [Grossberg, 2025; Kim, 2024], EEG-based BCIs [Singanamalla, 2022], seizure detection [Zhang, 2024], and motor imagery classification [Zhang, 2025]. Specialized frameworks such as NeuCube leverage spatiotemporal encoding and biologically inspired learning rules to achieve efficient EEG modeling and classification [Tan, 2020; Lim, 2025].

A key mechanism underlying biological SNNs is sparse coding, a strategy believed to support efficient sensory processing in cortical circuits [Barth, 2012]. Sparse activity patterns are shaped by excitatory– inhibitory balance, synaptic interactions, and cortical connectivity structures [King, 2013; Denève, 2016]. These principles motivate ongoing efforts to develop more biologically inspired computational models that incorporate temporal precision, stochasticity, and minimal reliance on global optimization.

Probabilistic spiking neural networks (PSNNs) extend classical SNNs by introducing stochastic elements that more closely mimic the variability of biological neurons. In PSNNs, spike generation and synaptic transmission occur according to probabilistic rules that capture uncertainty in neural responses [Jang, 2019; Nallathambi, 2021]. This stochasticity supports functions such as perceptual inference and decision-making [Pecevski, 2011; Deneve, 2004]. PSNNs also offer computational advantages. For example, unlike integrate-and-fire (I&F) neurons, which require continuous membrane-potential integration, probabilistic models rely on simplified spike-likelihood computations that scale well to large networks or neuromorphic hardware [Yao, 2025; Ocker, 2023].

In artificial neural systems, learning is typically driven by non-Hebbian rules such as backpropagation, which adjust synaptic weights based on global error signals rather than local spike coincidences [Gerstner, 2011; Panda, 2017; Kato, 2009; Jackson, 2020]. Although effective, these mechanisms are biologically implausible and do not resemble early stages of neural development. Before the maturation of activity-dependent Hebbian plasticity, synaptic organization in the developing brain is strongly shaped by activity-independent and synchronization-based processes, including structural remodeling, spontaneous activity, and coordinated bursts [Kerschensteiner, 2014; Kirchner, 2025; Matsumoto, 2024]. Neurons act as coincidence detectors: synchronous presynaptic spikes more effectively drive postsynaptic activation, improving signal-to-noise ratio and facilitating the establishment of functional circuits [Gansel, 2022; Sailamul, 2017; Brette, 2012; Chen, 2013; Buzsáki, 2012; Veit, 2023]. These synchronization-based principles provide a biologically grounded foundation for alternative learning rules that do not rely on backpropagation or Hebbian co-activation.

Motivated by this biological perspective, we propose a synchronization-based synaptic learning rule for probabilistic spiking neural networks. The rule is designed to emulate early developmental wiring mechanisms by strengthening connections based on coordinated spike timing rather than explicit supervisory signals or Hebbian correlations. The proposed model consists of a probabilistic three-layer feedback SNN architecture that incorporates two inhibitory layers functioning as probabilistic feedback pathways. Through the interaction of stochastic spiking and synchronization-based synaptic updates, the network autonomously self-organizes its inter-layer connectivity during training. This bio-inspired mechanism enables learning of spatiotemporal structure in EEG data without relying on backpropagation or labeled examples.

The remainder of this paper presents the proposed model, its learning dynamics, and preliminary results demonstrating its application to EEG-based decoding of auditory attention.

## Model & Methods

### EEG Data and Preprocessing

In this study, EEG data were collected from 16 participants engaged in an auditory attention task involving competing audio streams. The dataset is publicly available at Zenodo. For each participant, 24 minutes of EEG recordings corresponded to attention directed toward the left audio stream, and an additional 24 minutes corresponded to attention directed toward the right stream. Each participant’s recordings were segmented into 10 trials for left-ear attention and 10 trials for right-ear attention. Model training and testing were conducted on within-subject data to maintain consistency.

All EEG channels were re-referenced to the average of the mastoid electrodes. The signals were then band-pass filtered between 8 and 13 Hz and downsampled to 128 Hz. Each channel was normalized to zero mean and unit variance within each trial and scaled to the range [0, 1].

To convert continuous EEG signals into temporal spike trains, the **Threshold-Based Representation (TBR)** method was applied. In this approach, a spike is generated whenever the EEG amplitude exceeds a predefined threshold, with the spike timing corresponding to the threshold-crossing event. A range of threshold values from 0 to 1 was tested to identify the optimal parameter for analysis.

All simulations and analyses were performed in **MATLAB** (MathWorks Inc.), which served as the platform for neural network modeling, data processing, and visualization.

### Neural network’s architecture and spiking neuron model

#### Network architecture

A five-layer feedforward Probabilistic Spiking Neural Network (PSNN) was constructed, comprising both excitatory and inhibitory neuronal populations (Figure 1A). The input layer consisted of 64 spiking neurons, each representing one of the 64 EEG recording channels as temporal sequences. During training, preprocessed EEG time-series patterns were sequentially fed into this input layer.

**Fig. 1.**
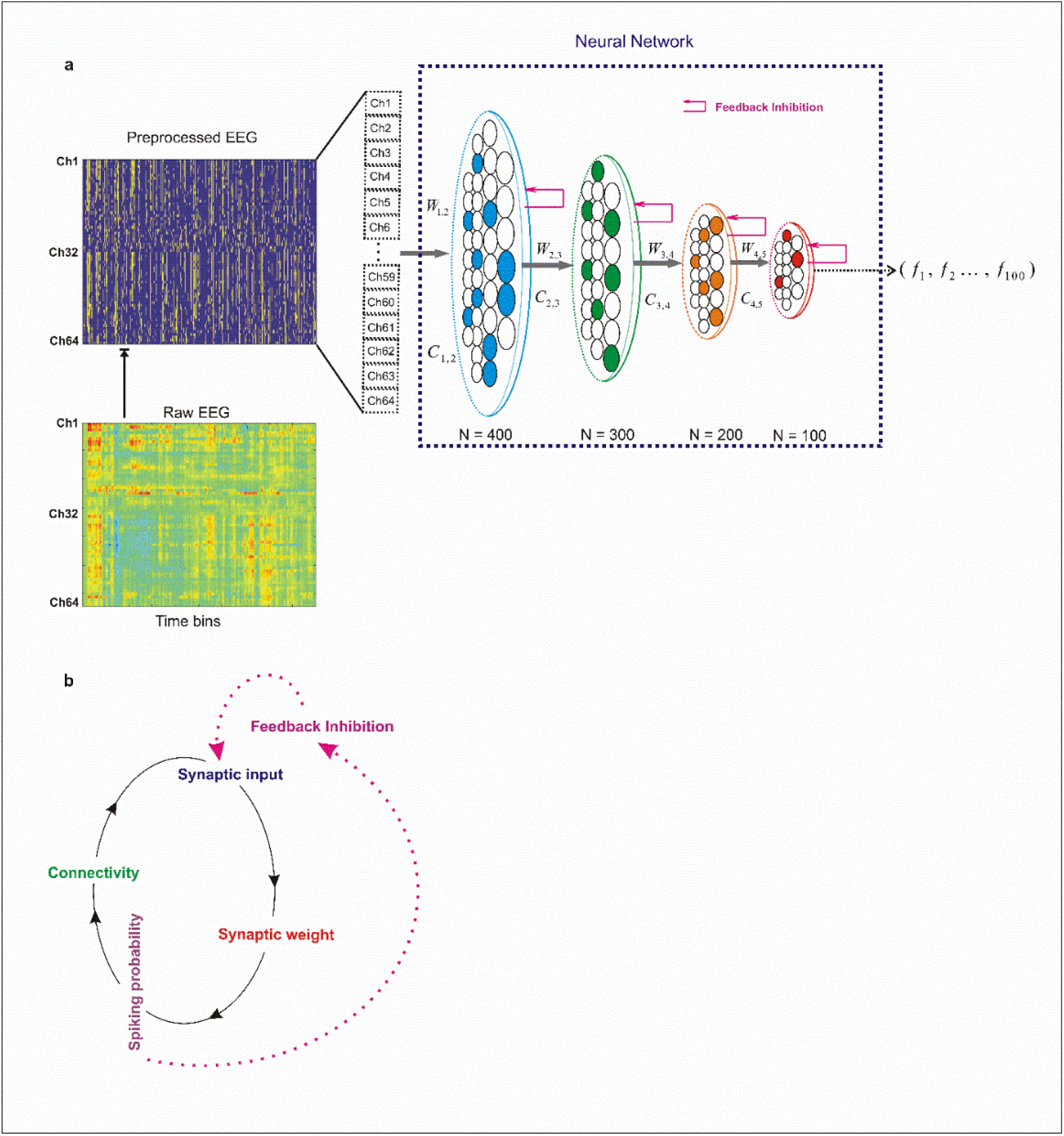
(a) Neural network architecture. The model comprises an input layer of 64 neurons, each corresponding to one EEG channel data. After preprocessing, these neurons generate spikes as were represented as binary events (“1”) within discrete time bins. These neural activities are presented into the network. The network includes four probabilistic excitatory spiking layers, each paired with a corresponding inhibitory layer in a fully connected feedback configuration. Every excitatory neuron is linked to a specific inhibitory counterpart, forming balanced excitation–inhibition pairs. All synaptic weights and connectivity values were initialized to 0.05 and updated dynamically during training. The activity of neurons in the fifth excitatory layer of the network (output layer) was analyzed to assess classification performance across simulations. **(b) Schematic of neural dynamics during training**. At the end of each training trial, synaptic weights are adjusted according to the temporal synchronization between presynaptic spikes. The spiking probability of each neuron depends on the balance of excitatory/inhibitory input, determined by both synaptic weights and incoming activity. Connectivity values are updated based on the postsynaptic neuron’s spiking probability, while feedback inhibition modulates overall synaptic drive through the inhibition-intensity parameter.

Subsequent layers were composed of probabilistic spiking neurons whose firing activity followed the network’s stochastic dynamics. Specifically, the second, third, fourth, and fifth layers contained 400, 300, 200, and 100 neurons, respectively. Each excitatory neuron was paired one-to-one with an inhibitory interneuron. Inhibitory neurons did not form mutual inhibitory connections; instead, each projected exclusively to its paired excitatory neuron, providing probabilistic feedback inhibition via recurrent synapses.

Synaptic efficacy between excitatory– inhibitory pairs was held constant at a weight of 1 throughout both training and simulation. At network initialization, the probability of forming synaptic connections between neurons in successive layers was set to 0.05, with initial synaptic weights sampled from a uniform distribution centered at 0.05. Both synaptic weights and inter-layer connectivity were subsequently adjusted during training via the network’s adaptive plasticity mechanisms. The spiking activity of neurons in the fourth layer was used to assess the model’s classification performance and overall discriminative capability.

#### Synaptic learning rule

In this study, we introduce a novel synaptic learning rule inspired by early neural development, during which neurons receive only a limited set of random synaptic inputs that are insufficient on their own to trigger action potentials. We hypothesize that the strengthening of synaptic connections in a postsynaptic neuron is determined by the degree of temporal synchronization among its afferent inputs. In this framework, synchronized presynaptic activity directed toward a postsynaptic neuron is assumed to convey meaningful information about the external environment.

Temporal synchrony among inputs is critical for shaping postsynaptic responses, as the precise timing of synaptic events can significantly influence neuronal excitability. When multiple presynaptic neurons release neurotransmitters simultaneously, or within a narrow temporal window, the postsynaptic neuron has an increased probability of firing, providing a robust mechanism for transmitting stimulus-related information. This synchrony allows neural networks to encode complex temporal patterns and supports higher-order processes such as sensory integration, selective attention, and memory formation. Moreover, neuronal synchronization contributes to rhythmic oscillatory activity in the brain, which is linked to large-scale functions including motor coordination, cognitive processing, and decision-making.

To formalize this non-Hebbian learning rule, we quantify the degree of synchronization among synapses converging onto a single postsynaptic neuron. Consider a population of *n* presynaptic neurons, each forming a single synaptic connection. These neurons fire stochastically according to their intrinsic firing probabilities, producing spike trains represented as binary vectors, where “1” denotes a spike and “0” denotes quiescence.

The proposed learning rule was implemented through the following steps:

##### Step 1: Computation of mean presynaptic activity

For each time bin, the mean spiking activity across all presynaptic neurons was calculated to estimate the overall firing level of the presynaptic population (Equation 1).

##### Step 2: Derivation of the synchronization score

The mean activity obtained in Step 1 was transformed into a synchronization score, which quantifies the degree of temporal coherence among the presynaptic inputs at each time bin (Equation 2).

##### Step 3: Weighted modulation of presynaptic spikes

At every time bin, the presynaptic spike vector was multiplied by its corresponding synchronization score (Equation 3), allowing individual spike contributions to be scaled according to the momentary level of network synchrony.

##### Step 4: Synaptic weight update

The average of these weighted spike values was then computed for each presynaptic neuron across the full trial duration (T), and the resulting value was used to update the corresponding synaptic weight (Equation 4).

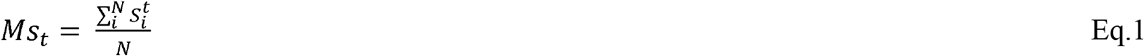

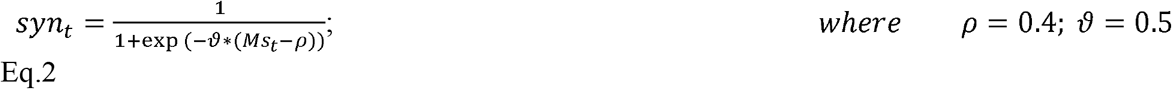

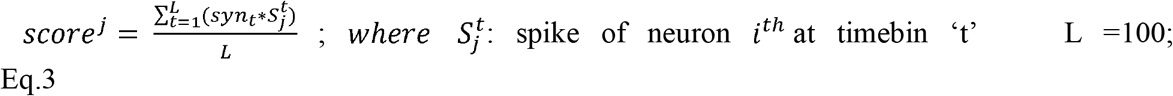

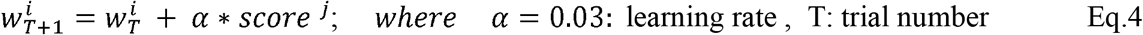

#### Neuron model and dynamics

The neuron model used in the proposed feedforward neural network is defined as follows. Neurons in the input (stimulus) layer are implemented as predefined probabilistic units, each initially connected to neurons in the second layer with a synaptic connection probability of 0.05. Neurons in the second and third layers are modeled as probabilistic spiking units, with their membrane potentials and firing probabilities determined by the synaptic inputs they receive, as described in Equation 5.

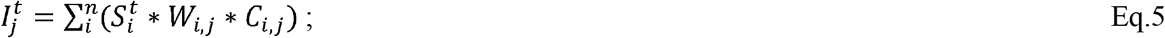

*n*: number of synaptic connections converging onto a single neuron

The spiking probability of each neuron at a given time bin is computed according to Equation 6. Synaptic connectivity between neurons and their preceding layer is updated at the end of each training trial following the rules defined in Equations 7–9. In these equations, *M*□ represents the average spiking probability of a neuron at the conclusion of a trial. Equation 8 is derived from prior research introducing a probabilistic model of structural plasticity, which captures the biological relationship between synaptic weight and synaptic volume, as well as the dependence of synaptic volume on synaptic lifetime.

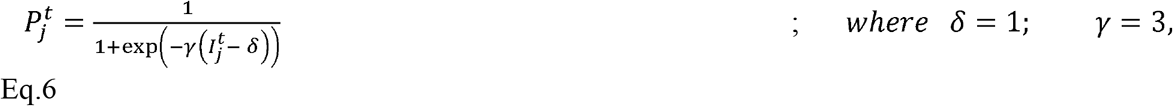

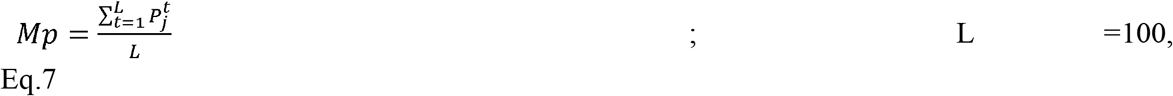

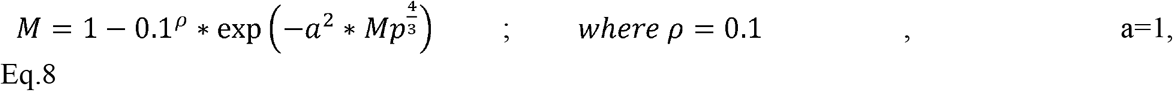

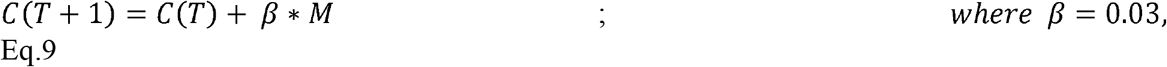

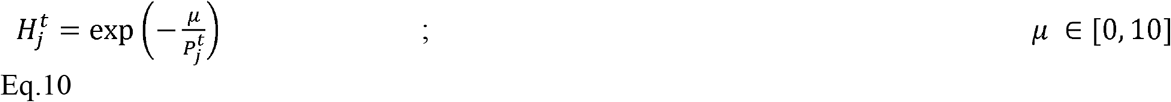

Within this framework, M represents the probability of forming new synapses and is used to calculate the updated connectivity of each neuron with its preceding layer. As training progresses, changes in connectivity alter the total synaptic input received by each neuron, which in turn affects its spiking probability, creating a closed-loop mechanism (Figure 1B).

To stabilize neuronal activity during training, feedback inhibition was incorporated. The activity of inhibitory neurons is modeled by Equation 10, where 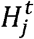 denotes the spiking probability of an inhibitory neuron receiving input from its paired excitatory neuron at time bin *t*. The parameter μ controls the strength of inhibition (Figure 5A). This inhibitory probability is then used to modulate the input activity of the corresponding excitatory neuron, providing a dynamic regulatory mechanism during learning. As a result, both neuronal connectivity and spiking probability are updated iteratively during training: connectivity is modified at the end of each trial, while feedback inhibition continuously modulates excitatory input in real time.

Thus, both the connectivity and spiking probability of each neuron are updated iteratively throughout training: connectivity is adjusted at the end of each trial, while feedback inhibition continuously regulates excitatory inputs in real time.

#### Network Training and classification accuracy

To train the network for EEG-based feature extraction, one sample from each of the leftward- and rightward-attention EEG datasets was selected and preprocessed. These training samples were presented to the input layer, which consisted of 64 neurons corresponding to the EEG channels. Two separate networks were trained independently for leftward- and rightward-attention EEG, respectively. After training, the firing rate patterns of the spiking neurons in the output layer of each network were recorded and stored for subsequent classification.

During testing, all EEG samples from both the left and right auditory streams were presented to the trained networks. Output layer firing rate patterns were computed and compared to the stored training patterns. The attentional direction of each test sample was assigned based on the network showing the highest pattern similarity, quantified using the Euclidean distance. Classification accuracy was calculated as the proportion of correctly classified samples out of the total 20 samples per subject.

#### Experiments

The classification system was systematically evaluated across a range of model parameters to determine the settings that maximize classification accuracy. In particular, multiple threshold values in the Threshold-Based Representation (TBR) method, ranging from 0.1 to 1, were tested. For each threshold, EEG signals were converted into spike trains, and the resulting network responses were used for classification. Performance was measured using classification accuracy, defined as the proportion of correctly classified samples across all trials for each subject. The threshold that produced the highest average accuracy across subjects was selected as the optimal TBR setting.

## Results

The goal of developing this deep spiking neural network is to extract discriminative features from EEG signals corresponding to rightward and leftward EEG data and represent them through the network’s connectivity patterns and synaptic weight configurations. To explore the dynamics of the proposed neural network model and determine the optimal conditions for classification accuracy, we systematically varied key parameters, with particular attention to the strength of feedback inhibition. Throughout the simulations, feedback inhibition proved crucial for maintaining network stability by constraining excessive synaptic growth and connectivity during training. Moreover, it substantially enhanced the model’s capacity to extract discriminative features from the EEG data.

**Figures 2 and 3** depict the temporal evolution of the average connectivity rate across network layers and the mean synaptic weights of neurons. The synaptic weights were initially initialized to 0.05. Following the presentation of EEG data corresponding to leftward and rightward stimulation—after appropriate preprocessing and conversion into spike-based activity—both measures exhibited dynamic modulation. The results obtained under three levels of feedback inhibition intensity (low, intermediate, and high) indicate that strong inhibition effectively suppresses the growth of both synaptic weights and layer connectivity. Conversely, weak inhibition leads to an uncontrolled increase in these parameters throughout the training phase. In contrast, an intermediate level of inhibition enables the model to maintain a balanced adaptive behavior, allowing for controlled growth of layer connectivity and synaptic weights, thereby promoting network stability and efficient learning dynamics.

**Fig. 2.**
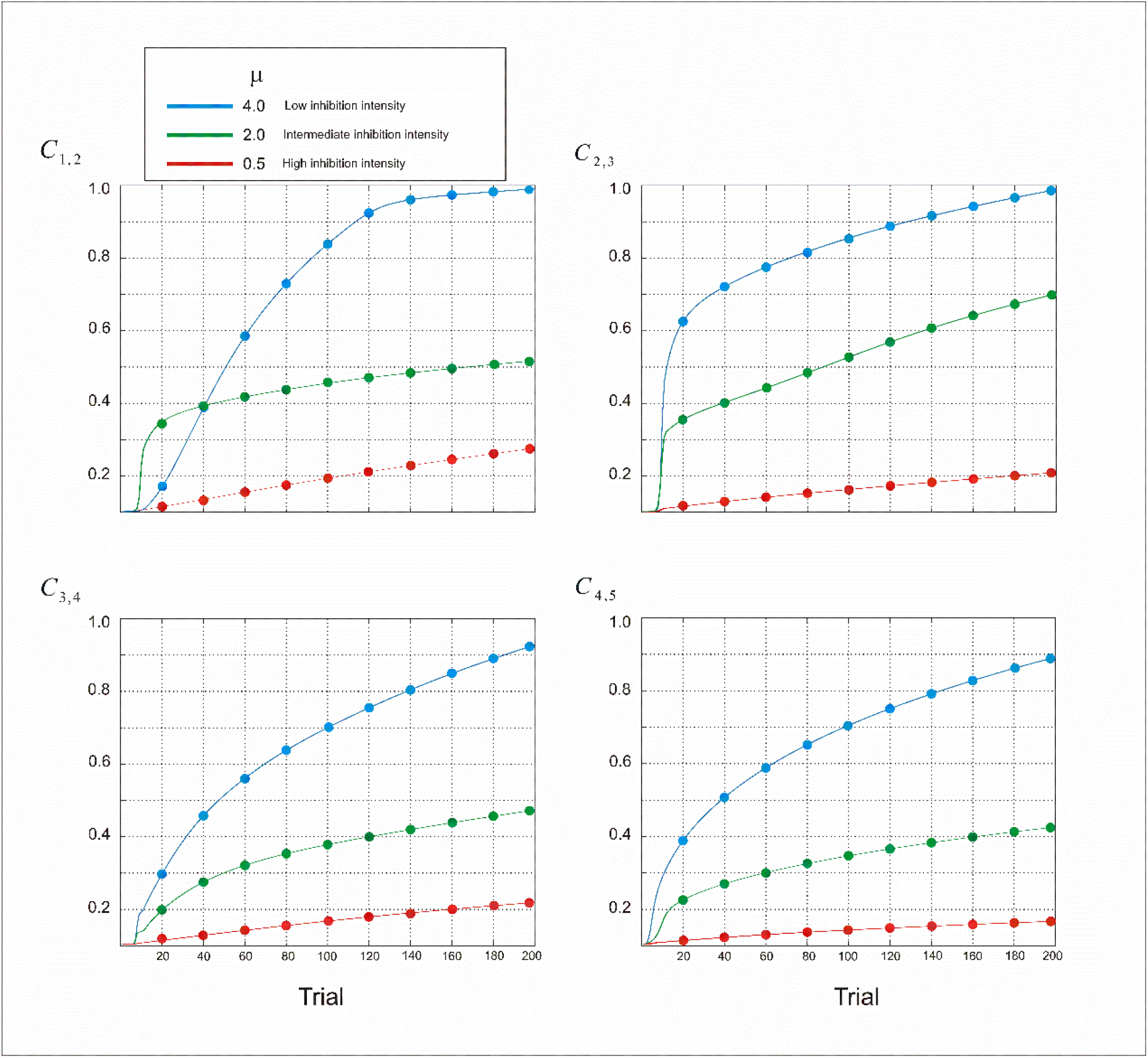
Dynamics of model parameters. Displayed are the temporal changes in mean connectivity rate of layers across network layers under varying levels of feedback inhibition. Simulation outcomes are shown for three inhibition intensities: low (μ = 4), intermediate (μ = 2), and high (μ = 0.5). Under low inhibition, connectivity rate exhibit uncontrolled growth over successive trials. In contrast, high inhibition suppresses connectivity expansion. The intermediate inhibition condition produces balanced dynamics, enabling stable yet adaptive increases in connectivity rate of the layers.

**Fig. 3.**
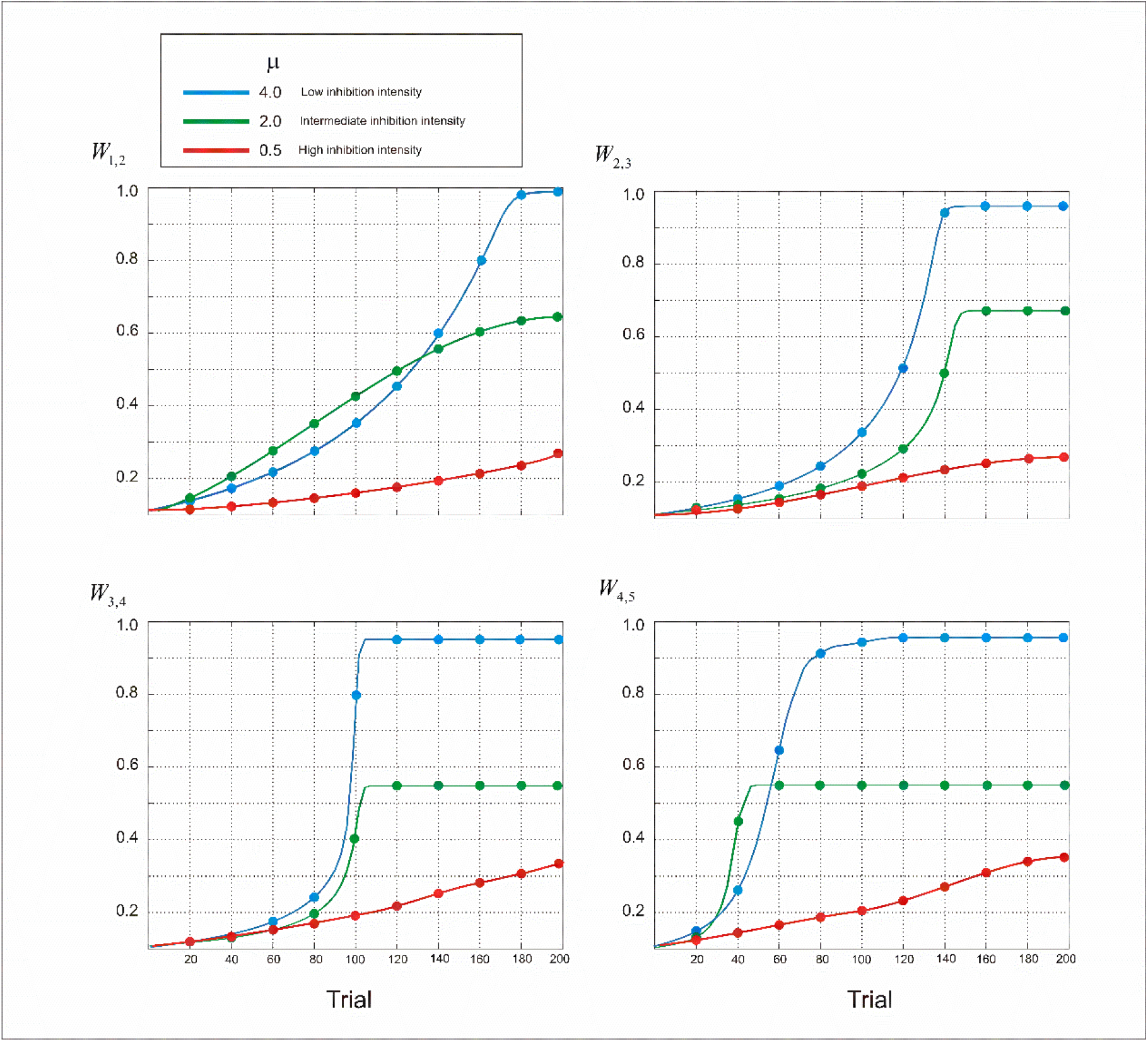
Dynamics of model parameters. Shown are changes in the mean synaptic weights under different levels of feedback inhibition. Simulation results are presented for three inhibition intensities: low (μ = 4), intermediate (μ = 2), and high (μ = 0.5). Low inhibition leads to uncontrolled growth of synaptic weights across trials, while high inhibition suppresses their increase. Intermediate inhibition achieves balanced dynamics, allowing controlled growth of synaptic strength of the neurons of layers.

**Figure 4**. shows the progression of the connectivity rate and synaptic weight development in the output-layer neurons of the model during training, in response to the presented EEG input data. The observed results indicate substantial heterogeneity in the activation patterns of individual neurons, suggesting diverse input processing dynamics. For this experiment, an intermediate feedback inhibition level was applied (μ =1). These findings underscore the critical role of the inhibition parameter in maintaining stable network dynamics. As shown in **Figure 5a**, inhibition probability varies with the parameter value and depends on the excitatory spiking probability. Specifically, lower parameter values result in stronger inhibitory effects, while higher values yield weaker inhibition of excitatory neurons across different activity levels. **Figure 5b** further demonstrates that as feedback inhibition intensity decreases, the excitatory spiking probability in the output layer increases correspondingly.

The simulations demonstrated that varying levels of feedback inhibition play a critical role in determining both the classification accuracy of the model and the connectivity rate between layers at the end of the training process. To explore this relationship further, we analyzed the connectivity rates of the layers alongside the corresponding classification accuracies throughout the training trials. **Figure 6a** highlights that sparse layer connectivity is essential for enabling efficient feature selection and accurate classification of input data. Furthermore, the model’s classification accuracy was found to be sensitive to the specific training samples drawn from the rightward and leftward EEG data. **Figure 6b** illustrates this dependency, showing how variations in the EEG data—coupled with noise—can influence the model’s performance.

**Fig. 4.**
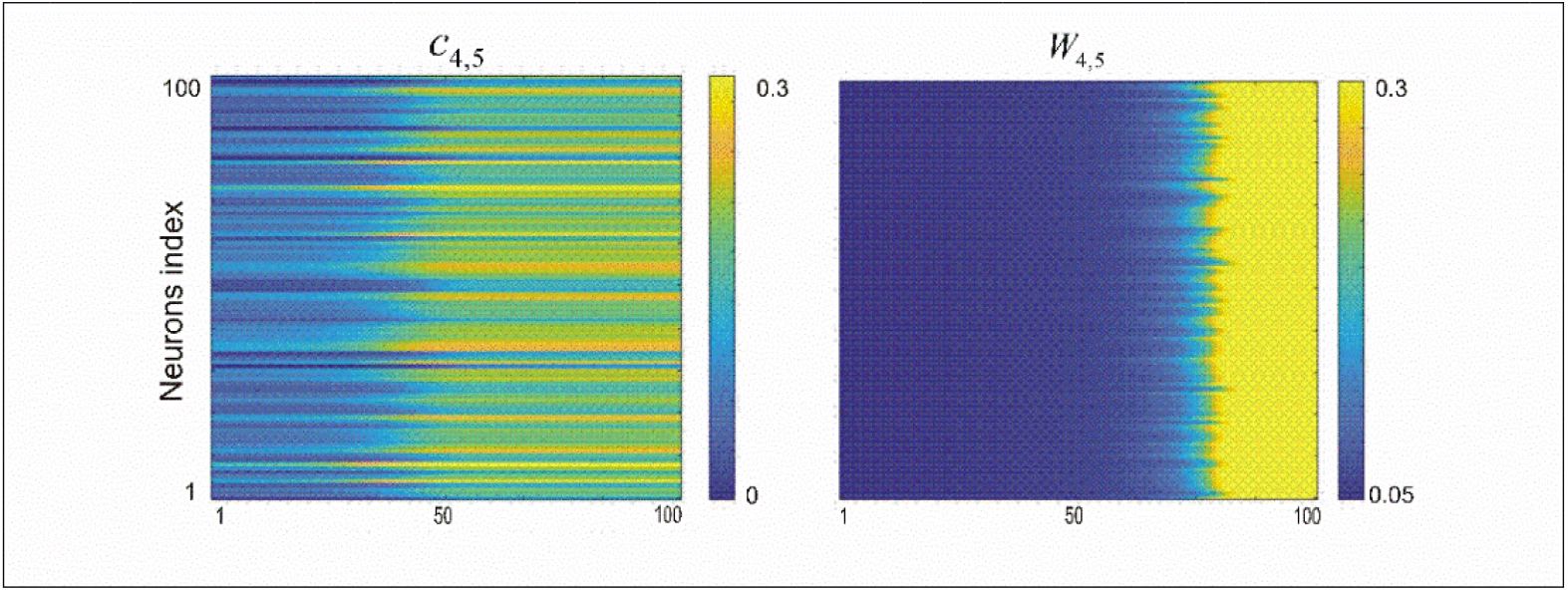
Evolution of model parameters under different inhibition intensities. The figure illustrates changes in synaptic weights and neuronal connectivity across training trials for an intermediate inhibition level (μ = 1). Distinct trajectories of synaptic adaptation and connectivity emerged among output-layer neurons, reflecting heterogeneity in their input activation patterns.

**Fig. 5.**
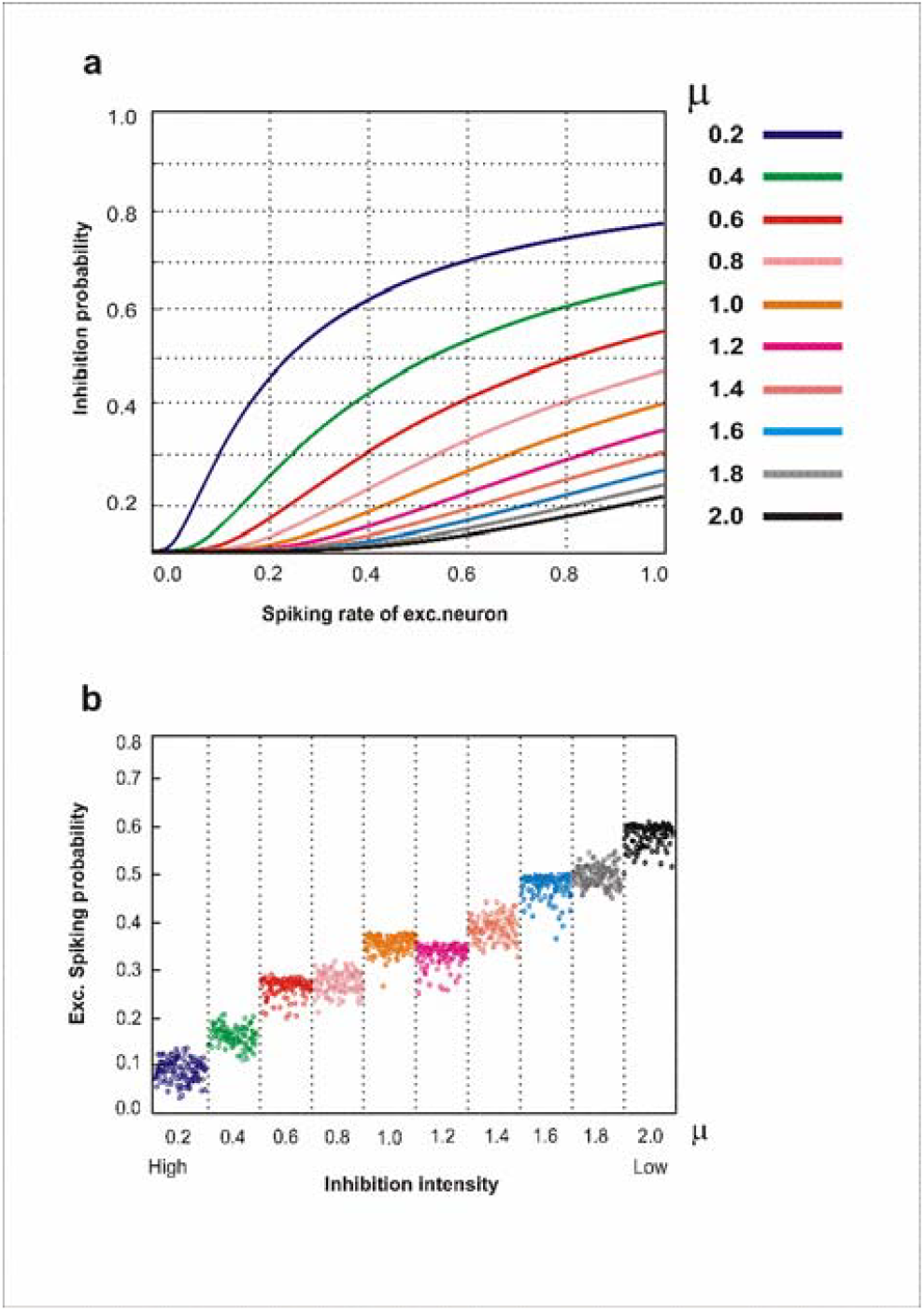
**(a)** Feedback inhibition probability of the inhibitory neurons as a function of excitatory spiking activity. Lower values of µ induce stronger inhibitory responses, whereas higher µ values attenuate the inhibition strength. **(b)** Excitatory spiking probability of output-layer neurons under varying levels of feedback inhibition. Elevated inhibition levels markedly suppress output-layer activity, while reduced inhibition intensity enhances the spiking probability of these neurons.

**Fig. 6.**
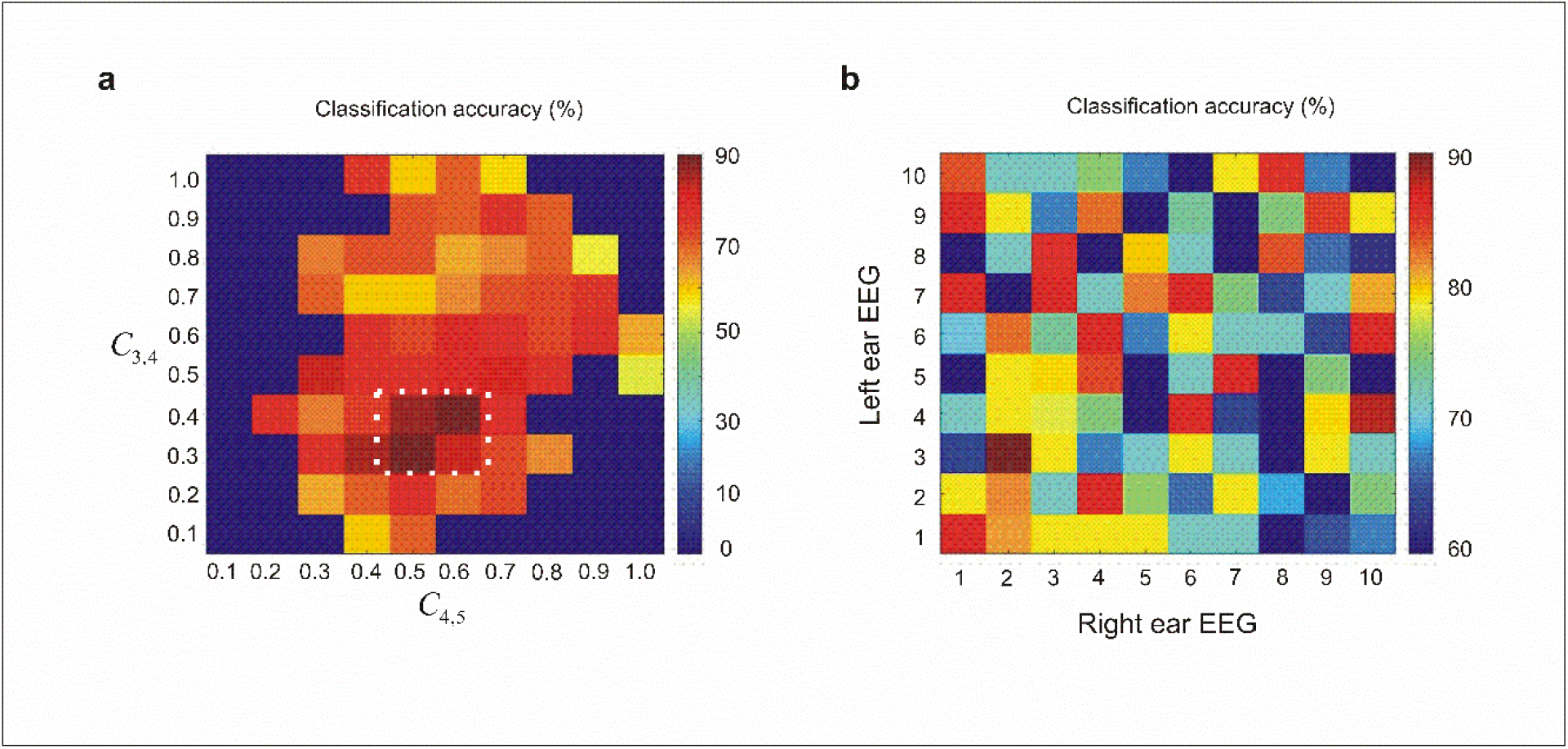
**(a)**. Influence of inter-layer connectivity on classification accuracy. The matrix presents the maximum classification accuracy across varying connectivity strengths between consecutive layers (*C*_3,4_ and *C*_4,5_). The optimal region, marked by the dotted white rectangle, varies with inhibition intensity, underscoring that sparse inter-layer coupling facilitates more efficient classification in feedforward spiking networks. **(b)** Dependence of classification performance on training sample selection. The matrix depicts classification accuracy for rightward and leftward attention samples, indexed by sample number. Variations in accuracy across sample pairs indicate that model performance is sensitive to the specific training inputs used. The experiments and simulations presented in the Results section correspond to the right-ear sample indexed as 10 and the left-ear sample indexed as 4 for one of the subjects.

The dynamics of synaptic synchronization are influenced by the model parameters (*θ*and *ρ*), which, in turn, affect both network behavior and classification performance.

**Figure 7a** shows the dependence of synaptic synchronization on the model parameters *ρ* and *θ*. Higher values of both parameters lead to stronger synchronization, with a nonlinear relationship observed between synchronization and these parameters. This reflects the complex interactions within the network, where increasing *ρ* and *θ* enhances synchronization for a given input level.

**Fig. 7.**
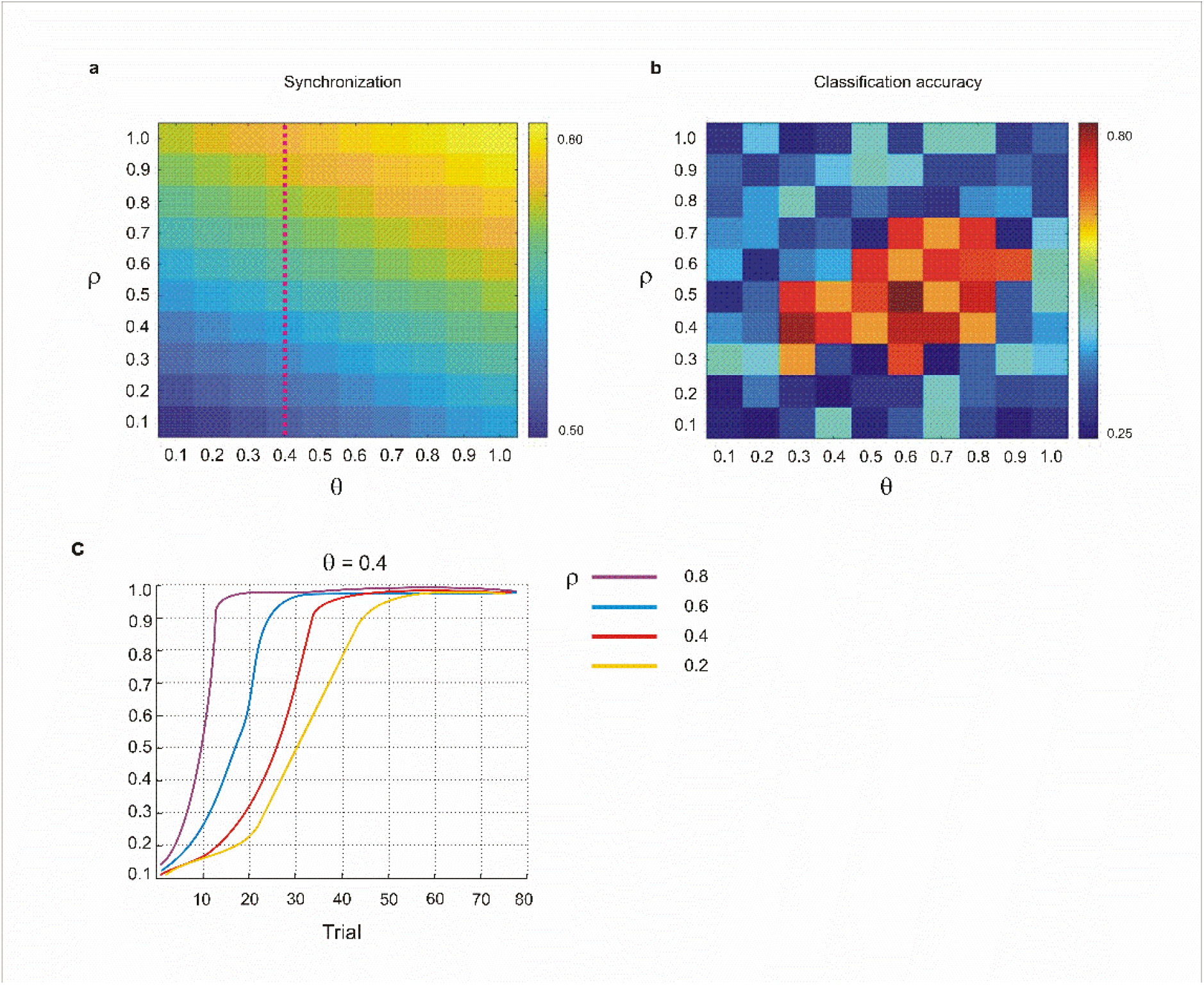
**(a)** Dependence of synaptic synchronization on model parameters *ρ* and *θ*. Higher values of *ρ*and *θ* lead to stronger synchronization for a fixed input level. Synaptic synchronization exhibits a nonlinear dependence on these parameters, reflecting complex interactions within the network dynamics. **(b)** Influence of synchronization parameters on synaptic weight evolution during training. For a fixed *θ* value (*θ* = 0.4; indicated by the red dotted vertical line in panel a) and four different ρ values, the evolution of synaptic weights in the output-layer neurons is shown. **(c)** Classification accuracy as a function of synchronization parameters, peaking at intermediate values under moderate feedback inhibition, suggesting an optimal regime for balanced information processing.

**Figure 7b** illustrates the impact of synchronization parameters on the evolution of synaptic weights during training. For a fixed *θ* value (*θ* = 0.4, indicated by the red dotted vertical line in panel a) and four different values of *ρ*, we observe how the synaptic weights in the output-layer neurons evolve over the course of training. This highlights the dynamic role of synchronization in modulating weight changes within the network.

Finally, **Figure 7c** shows the classification accuracy as a function of synchronization parameters. Notably, classification performance peaks at intermediate synchronization parameter values, with the best results occurring under moderate feedback inhibition. This suggests that there exists an optimal regime for balanced information processing, where both the network’s synchronization and inhibition levels are finely tuned for optimal performance.

In addition to the synchronization model’s parameters and feedback inhibition model’s parameter there exist some other parameters that influence the classification accuracy of the model.

In addition to the synchronization and feedback inhibition parameters, several other factors also impact the model’s classification accuracy. **Figure 8a** shows the classification accuracy across different feedback inhibition intensities, averaged over all subjects. Consistent with the previous findings from individual subjects, moderate levels of inhibition yield the highest accuracy following synaptic pruning during the training phase. In the absence of synaptic pruning, lower accuracies are observed across all inhibition intensities, underscoring the critical role of synaptic pruning in optimizing performance (see **Figure 8b**).

**Fig. 8.**
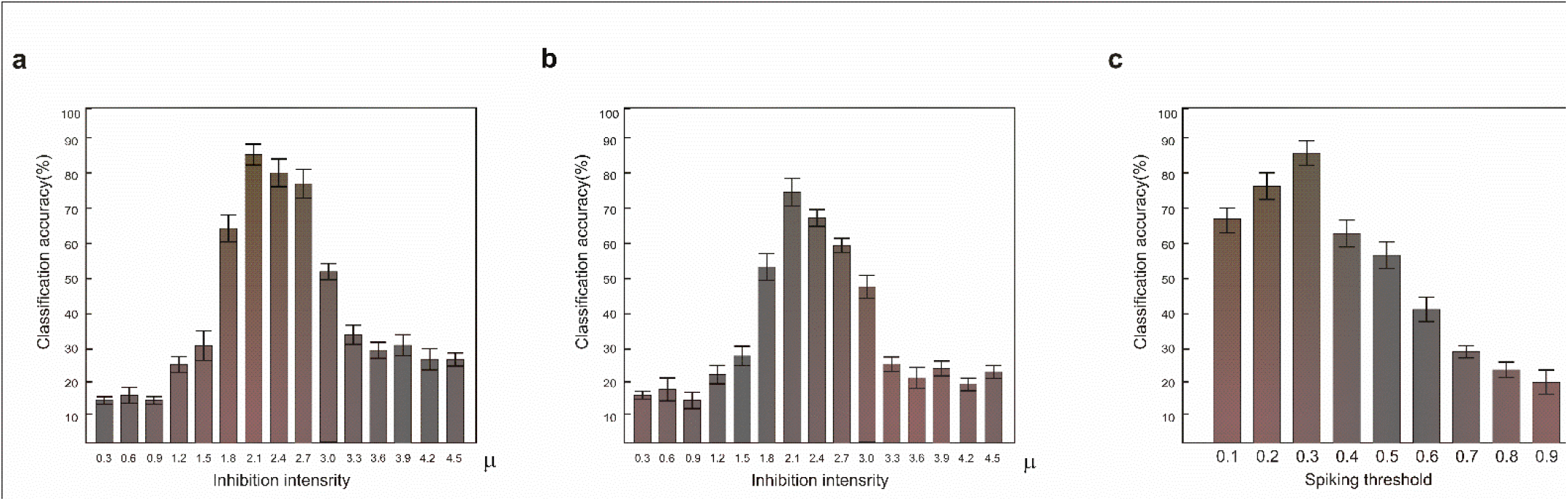
**(a)** Nonlinear dependence of classification accuracy on feedback inhibition intensity, with peak performance observed at intermediate inhibition levels. **(b)** Model classification accuracy across varying inhibition intensities in the absence of synaptic pruning. **(c)** Maximum classification accuracy as a function of input-layer spiking threshold, reaching its highest value at a threshold of 0.3.

**Figure 8c** examines the effect of the spiking threshold value applied during preprocessing of the input data on the model’s classification accuracy. The results indicate that the model achieves maximum accuracy when the threshold is set to 0.3.

## Discussion

The primary goal of this study was to develop a deep spiking neural network (SNN) capable of extracting discriminative features from EEG signals associated with rightward and leftward motor imagery tasks. Our findings demonstrate that the model’s performance, particularly its classification accuracy, is heavily influenced by a range of parameters, most notably feedback inhibition, synaptic synchronization, and preprocessing conditions. These results provide valuable insights into the dynamics of spiking neural networks and suggest directions for optimizing them for EEG-based classification tasks.

One of the most significant findings of this study is the crucial role of feedback inhibition in maintaining network stability and improving the model’s classification performance. Feedback inhibition governs the synaptic growth and connectivity of neurons during training, and our results, shown in **Figures 2** and **3**, reveal that an intermediate level of inhibition (μ = 1) leads to the most stable and efficient network dynamics. When inhibition was too weak, synaptic growth became unchecked, leading to excessive connectivity and overfitting, which compromised classification accuracy. Conversely, strong inhibition hindered the network’s ability to adapt and learn effectively by suppressing the growth of synaptic weights and connectivity too much.

These findings underscore the importance of tuning the inhibition parameter for optimal learning. The ability to balance synaptic growth while constraining excessive connectivity is key to ensuring that the network can extract meaningful features from the input data without becoming overly complex or prone to noise. This is especially important for EEG classification tasks, where the underlying signal is often noisy and variable across different subjects.

The temporal evolution of synaptic weights in response to EEG inputs, illustrated in **Figure 4**, highlights another important aspect of the model’s learning dynamics.

The observed heterogeneity in the activation patterns of individual neurons suggests that the network is capable of diverse input processing, a crucial feature for handling the complex, non-linear relationships inherent in EEG data. This diversity may help the model generalize better to unseen data, enhancing its robustness in classification tasks.

Interestingly, the intermediate level of feedback inhibition (μ = 1) allowed the model to maintain a balanced growth of synaptic weights, preventing both overfitting and underfitting. This finding is consistent with the broader understanding that neural networks—particularly spiking networks— require precise control over synaptic plasticity to achieve optimal learning. The heterogeneity observed also hints at the potential for selective feature extraction across different regions of the network, allowing the model to focus on the most informative aspects of the EEG signal.

Our results further show that sparse connectivity between layers is crucial for efficient feature selection and accurate classification of EEG data, as seen in **Figure 6a**. Sparse networks are often more efficient at identifying and processing the most important features of the input signal, as they avoid becoming bogged down by redundant or irrelevant information. This finding is particularly relevant in the context of EEG data, where the signal is high-dimensional and often contains much noise.

Moreover, the model’s sensitivity to specific training samples (e.g., rightward and leftward EEG data) underscores the importance of input variability and the model’s ability to generalize. **Figure 6b** demonstrates that variations in the EEG data, particularly due to noise, can significantly affect classification performance. This highlights a major challenge in working with real-world EEG data: ensuring that the model is robust to noise and able to generalize well across different subjects and conditions.

In addition to feedback inhibition, we also explored the impact of synaptic synchronization, governed by the parameters *ρ* and *θ*, on network dynamics and performance. As shown in **Figure 7a**, higher values of *ρ* and *θ* resulted in stronger synchronization, which enhanced the model’s ability to process the input data. However, the relationship between synchronization and network behavior is non-linear, suggesting that the model exhibits complex interactions between its parameters. **Figure 7b** further illustrates that synaptic synchronization modulates synaptic weight evolution, indicating that these parameters are not only important for network stability but also for controlling how the network adapts during training.

Interestingly, **Figure 7c** shows that classification accuracy peaks at intermediate synchronization values, particularly when combined with moderate feedback inhibition. This suggests that there is an optimal synchronization regime that balances the need for coordinated network activity with the ability to remain flexible and adaptive to input variations. These findings provide valuable insights into the trade-offs between synchronization and flexibility in spiking neural networks, which can guide future model design and parameter tuning.

The impact of preprocessing conditions on classification performance was also examined, particularly the effect of the spiking threshold applied during the conversion of EEG data into spike-based activity. **Figure 8c** indicates that setting the spiking threshold to 0.3 resulted in the highest classification accuracy. This finding highlights the importance of carefully selecting preprocessing parameters to optimize feature extraction. Given that EEG data are typically noisy and require preprocessing to convert continuous signals into spike trains, the choice of preprocessing settings can have a substantial effect on the network’s ability to learn meaningful patterns.

Our study demonstrates that spiking neural networks, when properly tuned, can effectively classify complex EEG signals. The findings suggest several avenues for future research. First, further exploration into the interaction between feedback inhibition and other network parameters, such as synaptic plasticity and learning rules, could provide deeper insights into how spiking networks learn to extract features from high-dimensional data like EEG signals. Second, the impact of network topology on performance warrants further investigation, particularly regarding how different layer connectivity patterns influence the efficiency of feature selection. Finally, incorporating more sophisticated preprocessing techniques and noise reduction strategies could further improve the robustness and generalizability of the model, particularly when applied to real-world EEG data from diverse subjects and tasks.

Sparse coding is a computational approach that represents data using only a small number of active features, focusing on capturing key patterns while minimizing redundancy. In deep learning models, sparse coding enhances efficiency and generalization by encouraging the network to learn compact, interpretable representations of data that is similar to how the brain encodes sensory information with a minimal number of active neurons. In this study, we demonstrate that sparse coding plays a pivotal role in improving the classification accuracy of deep learning models, particularly in spiking deep neural networks.

Conventional deep learning methods, such as convolutional neural networks (CNNs), typically require large training datasets. Our model requires only 10% of the data for training. This demonstrates the potential of the proposed probabilistic spiking neural network architecture for applications with limited training samples. Consequently, it may be of particular interest to researchers working with small datasets, such as EEG recordings.

In our earlier work, we implemented a feedforward spiking neural network for auditory attention classification and found that the maximum classification accuracy was achieved when the layers exhibited low connectivity rates. Building on this finding, the current study introduces a self-organized spiking neural network that dynamically determines optimal (sparse) inter-layer connectivity through the regulation of intermediate inhibitory intensity levels.

In conclusion, this study highlights the importance of key model parameters as feedback inhibition, synaptic synchronization, and preprocessing conditions in optimizing the performance of deep spiking neural networks for EEG classification. By carefully tuning these parameters, we were able to achieve stable network dynamics, effective feature extraction, and high classification accuracy. These findings contribute to the growing body of research on spiking neural networks and their potential applications in EEG-based brain-computer interfaces, where achieving both accuracy and efficiency is crucial. The insights gained from this work can inform the design of more advanced spiking neural networks for real-time, high-performance EEG analysis.

